# Blood Vessels Bioengineered from Induced Pluripotent Stem Cell Derived Mesenchymal Stem Cells and Functional Scaffolds

**DOI:** 10.1101/2025.02.03.636368

**Authors:** Luis Larrea Murillo, Zhongda Chen, Jun Song, Adam Mitchel, Steven Woods, Susan J. Kimber, Jiashen Li, Yi Li, Tao Wang

## Abstract

The development of small-diameter vascular grafts remains a major challenge in tissue engineering due to limited remodelling and regenerative capabilities. While strides have been made on the biofabrication of vessel mimics, little clinical translation success had been achieved to treat coronary artery disease (CAD). This study aimed to fabricate patient-specific bioengineered vessels using induced pluripotent stem cells (iPSCs) and functionalised biodegradable scaffolds. Human iPSCs were differentiated into mesenchymal stem cells (iMSCs) using SB431542, then further into vascular smooth muscle cells (VSMCs) with PDGF-BB and TGF-β1. Human bone marrow-derived MSCs (hBM-MSCs) were used to optimize differentiation protocols. Electrospun poly-L-lactide (PLLA) scaffolds coated with silk fibroin improved cell adhesion and proliferation. Both hBM-MSCs and iMSCs were seeded on these scaffolds for in-scaffold VSMC differentiation. The resulting cell-laden scaffolds were rolled into tubular structures (∼3 mm inner diameter, ∼20 mm length). Over 34–36 days, iPSCs differentiated into iMSCs expressing MSC markers (CD73, CD90, CD105), followed by successful VSMC differentiation within 9 days, confirmed by *α-SMA, CNN1, SM22*, and *MYH-11* expression. Silk fibroin-coated PLLA scaffolds enhanced MSC adhesion and proliferation compared to uncoated scaffolds. The engineered tubular grafts displayed VSMC markers and mechanical properties akin to autologous coronary artery bypass grafting (CABG) grafts. This study developed a versatile method to fabricate tissue-engineered blood vessels using stem cells and silk fibroin-coated scaffolds. The resulting grafts exhibited tunica media-like structures and mechanical properties comparable to autografts used in CABG, showing strong potential for clinical application.

## Introduction

Cardiovascular disease (CVD) remains the leading global cause of death, with coronary artery disease (CAD) responsible for more than 9 million fatalities each year ^1^. Revascularisation techniques have significantly improved the prognosis and quality of life for patients with vascular conditions, particularly in coronary artery disease (CAD) ^2^. Restoring coronary blood flow is essential for survival, and while percutaneous coronary intervention (PCI) has become more common as a less invasive non-surgical procedure, coronary artery bypass grafting (CABG) remains the preferred treatment for patients with extensive atherosclerosis, complex lesions, or comorbidities like diabetes ^3,4^. CABG requires vascular grafts to bypass occluded coronary arteries, with saphenous veins being the most commonly used autologous vessels ^5^. However, these grafts have a high failure rate, with 40-50% of grafts failing within 10 years due to intimal hyperplasia and associated complications. In contrast, arterial grafts such as the internal mammary artery and radial artery have better long-term patency (failure rates ≤10%), but require more invasive harvesting techniques and are prone to vasospasm ^6,7^. Additionally, many patients may not have suitable autologous vessels due to factors like frailty or advanced age, leaving up to 30% of CABG-eligible patients without an appropriate graft. This highlights the need for alternative graft materials ^8^.

Over the last two decades, tissue engineering has advanced significantly in developing vascular grafts that mimic the mechanical and physiological properties of native vessels. ^9,10^. To address the shortage of autologous grafts, acellular synthetic materials, such as Dacron and expanded polytetrafluoroethylene (ePTFE), have been clinically applied as alternatives, yet have fallen short in patency compared to autologous grafts, particularly in small-diameter applications ^11^. Small-diameter (<6 mm) tissue-engineered vascular grafts (TEVGs) have demonstrated success as arteriovenous shunts in hemodialysis patients, but their application in coronary artery bypass remains limited ^12^. Decellularized allogenic saphenous veins repopulated with autologous endothelial cells (ECs) have reached the clinical setting for CAD treatment, but with a low degree of success ^13^.

An essential aspect of vascular tissue engineering is the scaffold, which provides structural support for growing tissue. Electrospinning, a low-cost and versatile method, has been used to create fibrous meshes from synthetic polymers like poly(L-lactic acid) (PLLA) or polycaprolactone (PCL) and has demonstrated to be a good candidate for fabricating scaffolds for vascular grafts ^14,15^. While several electrospun TEVGs have shown promising results *in vivo*, none have reached the clinic to treat CAD. This has been attributed to elastic modulus differences observed in electrospun TEVGs which greatly exceed those observed in native vessels and have been identified as a significant cause of graft failure ^16,17^. To improve biocompatibility, electrospun scaffolds have been coated with natural biomaterials such as silk fibroin. Silk fibroin coatings have shown to not only improve cell function due to the cell adhesive motifs it possesses but degradation can be easily tunned for optimal tissue growth and formation ^18^. Furthermore, modifications to surface topography have also shown to further enhance biocompatibility of these scaffolds as porous/rougher surfaces can improve cell attachment and migration of various vascular cell types ^19,20^.

Another key aspect in vascular tissue engineering lies in sourcing appropriate cells for graft construction. Currently, most TEVGs that have reached clinical trials have been derived from autologous primary vascular cells such as fibroblasts, vascular smooth muscle cells (VSMCs), and ECs. However, these primary cells have limited regenerative capacity due to the donor’s age, with poor proliferative ability and a high risk of senescence. Moreover, harvesting and culturing these cells is both time-consuming and expensive ^16,21^. These challenges have driven interest in stem cell technologies, particularly the use of induced pluripotent stem cells (iPSCs) and mesenchymal stem cells (MSCs). IPSCs offer an attractive solution due to their self-renewal capacity and pluripotency, akin to embryonic stem cells (ESCs) but without associated ethical concerns ^21,22^. MSCs, although more limited in differentiation potential, exhibit excellent genomic stability and have well-established clinical safety. Additionally, MSCs have immunosuppressive properties, making them suitable for both autologous and allogenic therapies ^22,23^.

In this study, we present a viable method to fabricate tissue engineered blood vessels by seeding human iPSCs (hiPSCs)-derived MSCs (iMSCs) onto silk fibroin fuctionalised biodegradable electrospun scaffolds and allowing in-situ differentiation of VSMCs. The fabricated vessels exhibited tunica media-like features and mechanical properties similar to autologous CABG grafts. Their simplicity suggests potential as an “off-the-shelf” solution.

## Materials and Methods

### Culture of iPSCs

E1PL1 (donor <u>SAMEA2536417</u>), SERU7 (donor <u>SAMEA3965165</u>) iPSC lines were established from human dermal fibroblasts using a CytoTune iPS Prprogramming kit (Life Technologies) by HipSci (Wellcome Genome Campus, UK) and made available via Public Health England through European Collection of Cell Cultures (ECACC) under a material transfer agreement. The SW171A wildtype hiPSC line was provided by Professor Kimber’s laboratory (University of Manchester, UK) and derived as previously described ^24^.The iPSCs were maintained in Essential 8™ Medium (Gibco, A1517001, USA) and seeded on Matrigel coated 6-well plates. Once thawed, they were passaged when 70-80% confluency was reached. Cells were passaged at least two times by dissociating with 0.5 µM EDTA (ThermoFisher, 15575020, USA) to remove any differentiated cells present after thaw prior to conducting characterisation and differentiation experiments. All cell-lines were maintained at 37°C with 5% CO2 incubation conditions.

### Culture of hBM-MSCs

Human bone marrow derived mesenchymal stem cells (hBM-MSC) (C-12974) harvested from normal human bone marrow from individual donors were cultured with Mesenchymal Stem Cell Growth Medium 2 (C-28009) purchased from PromoCell (Germany). Culture medium was changed every 2-3 days and passaged once ∼90% confluence was reached for further expansion or experimental use. All experiments with hBM-MSC were conducted between passages 3 to 7.

### Generation of iMSCs from iPSCs

Generation of hiPSC derived MSCs (iMSCs) were obtained via a monolayer-directed approach by Chen et al. ^25^ with significant modifications. Briefly, hiPSCs were seeded on Matrigel (Corning, 354230, USA) coated plates at a 25,000 cells/cm^2^ density and cultured with a 10uM ROCK inhibitor supplemented Essential 8™ Medium for the first 16-24 hrs. Thereafter, ROCK inhibitor was removed from the Essential 8™ Medium and maintained for another 48-72 hrs until a ∼70-80% confluence was reached. To initiate differentiation (Day 0) medium was changed to a Dulbecco’s Modified Eagle’s medium F-12 (DMEM F-12) by Gibco (11320-074, USA) supplemented with 1mM L-Glutamine, 1% (x1) MEM Non-essential Amino Acid Solution, 1% Penicillin-Streptomycin, 10µM, SB431542 in DMSO and 20% knockout serum replacement (KOSR). After 10 days, cells were disassociated with TrypLE Express (Gibco, 12604021, USA), resuspended on Mesenchymal Stem Cell Growth Medium 2 (Promocell, C-28009, Heidelberg, Germany) and seeded on uncoated plates at a density of 80,000 cells/cm^2^. When cells reached 80-90% confluence, they were first passaged at a density of 120,000 cells per cm^2^. Thereafter, cells were passaged at 1:3 or 1:2 ratios for the remaining of the culture period. At days 34-36, iMSCs positive for common MSCs markers were generated and maintained with the Mesenchymal Stem Cell Growth Medium 2.

### Osteogenic differentiation of hBM-MSCs and iMSCs

HBM-MSCs and iMSCs maintained in Mesenchymal Stem Cell Growth Medium 2 were dissociated with TrypLE Express once ∼ 90% confluence and seeded on 10 µg/ml human fibronectin (Promocell, C-43050, Heidelberg, Germany) coated plates at a density of ∼120,000 cells/well. When cells reached a 100% confluence, culture medium was changed to the Mesenchymal Stem Cell Osteogenic Differentiation Medium (C-28013) by PromoCell (Heidelberg, Germany) to induce osteogenic differentiation. Osteogenic differentiation medium over a 28-day differentiation period. At day 28, Cells were fixed and stained with Alizarin Red S to detect extracellular calcium deposits and mineralisation in osteogenic differentiated cells.

### Generation of VSMCs from hBM-MSCs and iMSCs

HBM-MSC derived VSMCs (hBM-MSC-VSMCs) and iMSC derived VSMCs (iMSC-VSMCs) were generated using the following protocol. HBM-MSCs or iMSCs resuspended on Mesenchymal Stem Cell Growth Medium 2 were seeded on plates at a density of ∼300,000 cells/well. After allowing cells to attach overnight, Medium was changed to VSMC induction medium consisting of Dulbecco’s Modified Eagle’s medium (DMEM) (Sigma-Aldrich, D6429-500ML, United Kingdom), 10% fetal bovine serum (FBS) and 1% Penicillin-Streptomycin. Medium was further supplemented with Transforming growth factor beta 1 (TGF-β1) (HEK293 derived) and Platelet-derived growth factor (PDGF-BB) from PeproTech (Rocky Hill, United States) at a concentration of 2ng/ml each. Cells were cultured under the differentiation medium for 9 days with medium being changed every 2-3 days.

### Immunocytochemistry

Cells were fixed with 4% paraformaldehyde (PFA) and permeabilised with 0.1% triton X-100 in PBS for detecting intracellular makers. Subsequently, cells were incubated with primary antibodies overnight at 4°C and secondary antibodies for 1 hr at room temperature thereafter. To determine successful differentiation of iMSCs, cells were stained with common MSC membrane markers CD44, CD73, CD90 and CD105. Generated VSMCs from either hBM-MSC or iMSCs were stained with intracellular markers α-SMA, CNN1, SM22 to determine its successful differentiation. Furthermore, hBM-MSC-VSMCs and iMSCs-VSMCs we stained with CACNA1C to assess their contractile function. Finally, all cells were counterstained with DAPI (Sigma-Aldrich, D9542, USA) to stain the nucleus and cells assessed for morphology had actin filaments stained with Phalloidin-iFluor 488 (Abcam, ab176753, UK).

### Flow Cytometry

To prepare samples for flow cytometry, cells were disassociated with acutase (Thermofisher, 00-4555, USA) and suspensions were first stained with fixable viability dye eFluor 780 (eBioscience, 65-0865-14, USA) for 30 minutes at 4°C. Thereafter, cells were fixed with 4% PFA and stained with antibody against cell surface protein markers in accordance with manufacturers recommendations. For intracellular markers, permiabilisation was conducted with 2% FBS in PBS and 0.1% Triton X-100 for 20 minutes before incubation with antibodies. Compensations were performed using unstained cells, viability dye stained cells and UltraComp ebeads by eBioscience (San Diego, CA, USA) conjugated with each antibody. Compensation and flow cytometry experiments and analysis was performed on the LSRFortessa SORP cytometer (BD Biosciences, USA). Cells were stained for pluripotent markers SSEA4, TRA-1-60 and Oct 3/4. Differentiating cells were also stained for positive MSC makers CD73, CD90 and CD105 as well as negative MSC markers HLA-DR, CD34 and CD14 to determine their multipotency

### Quantitative Real-Time Reverse Transcription-Polymerase Chain Reaction

Quantitative Reverse Transcription Polymerase Chain Reaction (qRT-PCR) was performed on hiPSC to MSC and iMSC/hBM-MSC to VSMC differentiating cells at different time-points. Total RNA extraction was performed using the RNeasy mini kit by Qiagen (Venlo, Netherlands) according to the manufacturer’s instructions. Reverse transcription for cDNA synthesis was performed using the High-Capacity RNA-to-cDNA™ Kit (4387406, Thermofisher, USA). The cDNA was added to a master mix composed of PowerUp™ SYBR™, primers for specific genes assessed and DEPC-Treated water. Mix was aliquoted (20 μl) into each well of a 96-well plate and transferred onto the StepOne™ Real-Time PCR System to run a Comparative Ct (ΔΔCt) experiment. House-keeping gene Glyceraldehyde 3-phosphate dehydrogenase (GAPDH) and genes of interest were evaluated using the 2^-ΔΔCT^ to analyse fold gene expression changes amongst the samples.

### Fabrication of electrospun scaffolds and functionalisation

Electrospun scaffolds were fabricated using a protocol previously described ^19^. Briefly, 2 wt% Poly-L-lactide acid (PLLA, PL 65) in Dichloromethane (DCM):N,N-Dimethylformamide (DMF) (19:1) solutions were loaded onto Luer plastic syringes with 21-gauge needles. Loaded syringes were placed on a nanofibre electrospinning unit (TL-Pro, Tongli Tech, China) and PLLA solutions were expelled at a controller flow rate of 5 mL/h. Fibres were spun using a voltage supply of 23 kV and at a walking distance of 30 cm from the collectors. After 4 hrs, Pristine fibrous scaffolds were removed from the collector and immersed in ≥ 99.8% acetone for 5 min to produce porous fibre scaffolds. Scaffolds were then further functionalised by covering the surface with a 1 wt% silk fibroin solution diluted in ethanol (EtOH) and dH_2_0 (Silk fibroin:EtOH / 2:1). Scaffolds were left to air dry after adding the silk fibroin solutions and via cross-linking inactions between EtOH and silk fibroin, a solid silk fibroin coat was left on the surface of fibrous scaffolds.

### Seeding hBM-MSC and iMSCs on Electrospun Scaffolds

Pre-seeding, scaffolds were pre-conditioned by incubating the with 10% FBS supplemented DMEM by incubating at standard cell culture conditions of 37°C 5% CO^2^ for 4-6 hrs to aid cell attachment. Thereafter, when hBM-MSC and iMSCs reached ∼90% confluence, cells were re-suspended and seeded on ∼15 mm diameter scaffolds at a density of 150,000 cells/scaffold. Cells were cultured with Mesenchymal Stem Cell Growth Medium 2 during the first 24 hours of attachment and subsequently switched to the VSMC differentiation medium detailed above for the next 9 days. Thereafter, medium was switched to a 10% FBS supplemented DMEM maintenance medium for the Remainer of the time depending on the experiment with medium changed every 2-3 days.

### Alama Blue Assay

Cells seeded on ∼15 mm diameter scaffolds were used to assess the biocompatibility of pristine, porous and silk fibroin coated scaffolds. The metabolic activity of cells laden on scaffolds was measured using an AlamarBlue™ (Invitrogen, DAL1025, USA) assay. Absorbance of AlamarBlue™ at 570 nm, using 600 nm as a reference wavelength (normalized to the 600 nm value) was measured using the FLUOstar® Omega microplate reader by BMG LABTECH (Offenburg, Germany). Reduction of AlamarBlue™ was calculated to determine metabolic activity of cells. To assess proliferation, cells were disassociated with TrypLE Express, resuspended and stained with Trypan Blue Solution, 0.4% (Gibco, 15250061, USA). Cell susoensions were then placed on Cell Counting Chamber Slides and inserted on a Countess II FL Automated Cell Counter (Invitrogen, USA) to measure changes in cell numbers throughput culture. AlamarBlue™ and cell count measurements were taken at different time-points (Day 1, 4, 7 and 10) to assess metabolic activity and proliferation of cell-laden scaffolds.

### Fabrication of Tissue Engineered Blood Vessels

HBM-MSC and iMSCs were seeded on 2.5 x 5.5 cm silk fibroin coated sheets using the same cell seeding protocol detailed above at a density of 150,000/cm^2^. After the 9-day VSMC induction period, hBM-MSC-VSMCs and iMSCs-VSMCs were cultured with the 10% FBS supplemented DMEM maintenance medium for 21 days (3 weeks total). After 3 weeks, the cell-laden sheets were wrapped around a 3 mm diameter stainless steel rod and from a tubular structure. For the tube to maintain its shape, an alginate sealing was applied using a modified rapid-casting method published by Ghanizadeh Tabriz et al. ^26^. Briefly, the cell-laden sheets were wrapped around the rod were dipped in 4% (w/v) alginate solutions for 3 sec. Subsequently, alginate coated constructs were dipped in a 100mM CaCl2 cross-linking solution for 3 min. Thereafter, the construct with the solidified alginate seal was removed from the rod leaving a hollow space in the middle of the cylindrical structure with similar dimensions to that of a small-diameter artery. Constructs using silk fibroin coated sheets without cells were also produced via the same method for mechanical analysis comparisons.

### Mechanical Properties Assessment of Tubular Constructs

To assess the mechanical properties of Tubular constructs, a tensile test experiment was conducted. The mechanical tensile test was performed on constructs approximately 20 mm in length, 3 mm in inner diameter and 4 mm of outer diameter with an Instron 3344 (MA, USA) static tensile test equipped. Tubes were connected distally and proximally onto plugs with flat ends that were fixed on the upper and lower grips of the instrument. The Instron instrument ran with a 10 Newton cell load and a deforming rate of 5 mm/min to extend tube constructs until fracture. Data collected to measure the ultimate tensile strength (UTS) was used calculate the theoretical burst pressure (P) using the following equation where (T) represents the UTS, (do) the outer diameter and (di) the inner diameter of constructs.

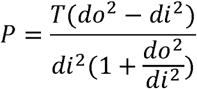

### Statistical Analysis

Statistical analysis was conducted and values were analysed using the GraphPad Prism software and Microsoft Excel 2016. One-way ANOVA test and a Tukey’s multiple comparisons test was applied to assess for statistical differences in experiments containing more than two groups. When p-values were <0.05, differences were considered significant. Data is presented as mean ± standard mean error (SEM).

## Results

### Producing iMSCs from iPSCs

To take the advantages of iPSCs that are able to provide inexhaustible source of autologous MSCs that have great regeneration potential and are easy to culture and differentiate into vascular cell types, we first established a robust protocol to produce MSCs from iPSCs. This protocol was based on Chen et al. ^25^ with modifications as illustrated in **Figure 1A**. Prior to differentiation, the pluripotency of hiPSCs was re-confirmed for hiPSC markers including SOX2, SEEA4, Oct3/4 and TRA1-60 via immunofluorescence staining and flow cytometry (**Figure S1A-B**). During differentiation, the expression of *NT5E* (CD73) and *ENG (*CD105) and *ENG (CD105)* steadily increased during the 34 days of iMSC differentiation as determined by RT-qPCR. By day 34, the expression of *NT5E* (CD73) and *ENG (CD105)* increased ∼170-fold and ∼26-fold, respectively, compared to the start of hiPSC differentiation (day 0) (**Figure 1B**). This was further validated by the flow cytometry data showing that the percentage of cells positive for CD73 and CD105 increased from 12.7% and 0.25% on day 0 to 75.55% and 50.75% on day 36, respectively (**Figure C**). The MSC marker CD90, in the other hand, did not increase further from the iPSCs during the differentiation (**Figure 1B and C**), however, the percentage of iMSCs positive for CD90 were comparable to those observed in hBM-MSCs (92.60%). The increase in MSC markers was also accompanied with a decrease in of pluripotent factors SEEA4 and TRA-1-60 (**Figure S2**). Moreover, iMSCs lacked key MSC negative marker CD34 and low levels for CD14 and HLA-DR, which were also similar to those of hBM-MSCs (**Figure S3**). Lastly, differentiated iMSC were observed to display a spindle-shaped and elongated morphology as well as positively exhibited MSC markers including CD44, CD105, CD73 and CD90 by immunofluorescent staining. Morphology and positive markers were comparable to those of hBM-MSCs (**Figure 1D**).

**Figure 1.**
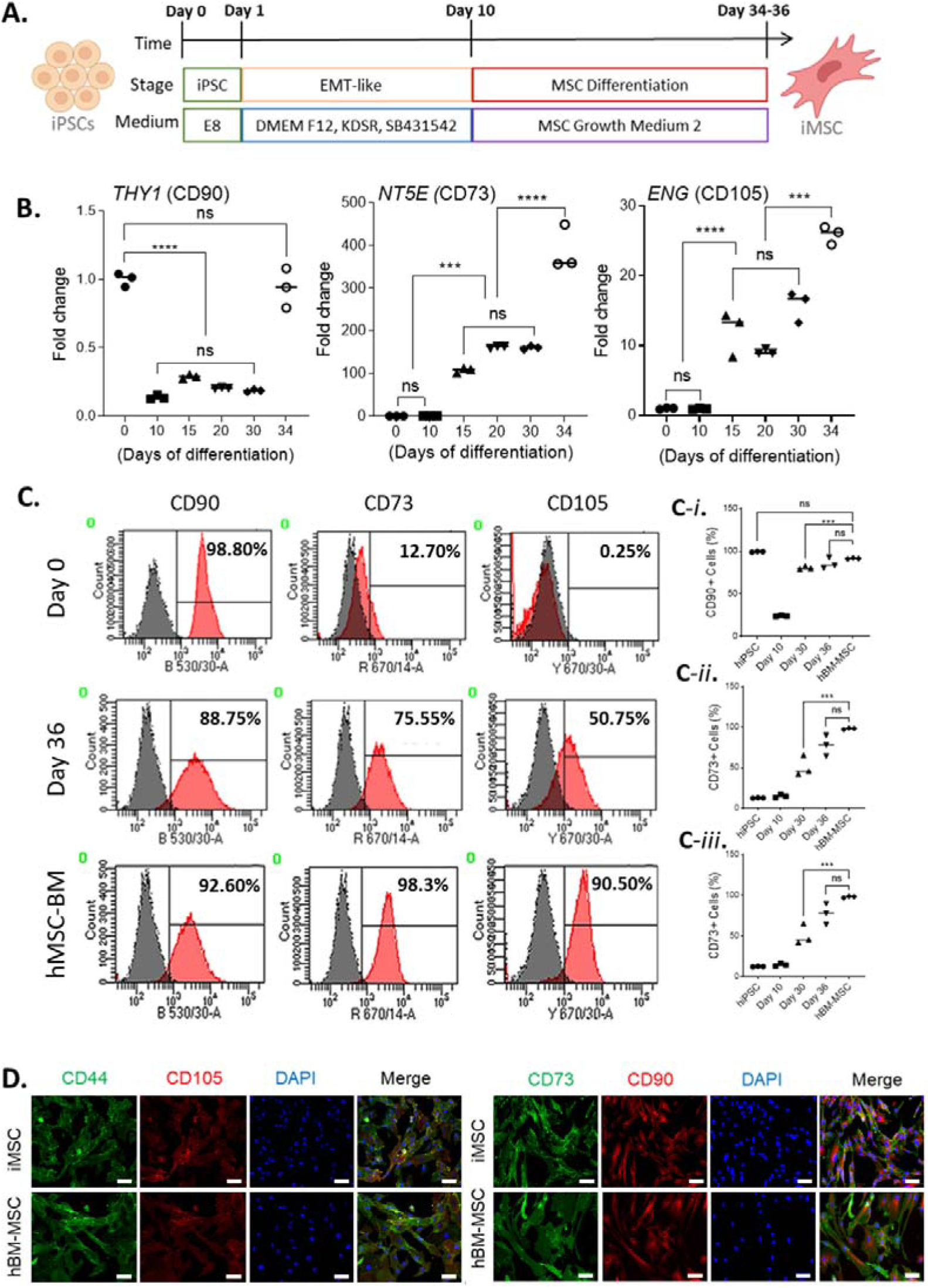
hiPSC differentiate into cells with an MSC phenotype exhibiting common MSC markers. A schematic of the MSC differentiation protocol is shown (**A**). Fold gene expression increases in MSC marker-genes *THY1 (CD90)*, *NT5E (CD73)* and *ENG (CD105)* was observed throughout differentiation of hiPSC-iMSCs (**B**). Histograms for common MSC positive markers are displayed (**C**) and increase of the percentage of cells positive for CD90, CD73 and CD105 after 36 days could also be noticed via flow cytometry (**C-i to C-iii**) after 36 days when iMSCs were derived. Data significance is presented as *** P ≤ 0.001 and **** p ≤ 0.0001 (n = 3).

iMSCs were then functionally verified by their ability to differentiate into osteogenic cells, a hallmark characteristic of the multipotency for MSCs. The iMSCs derived from 3 independent iPSC lines proved to be able to undergo osteogenic differentiation as evidenced by Alizarin Red S staining 28 days after differentiation (Figure 2 a-i), similarly to that of the parallel control hBM-MSCs (Figure 2 j-l).

**Figure 2.**
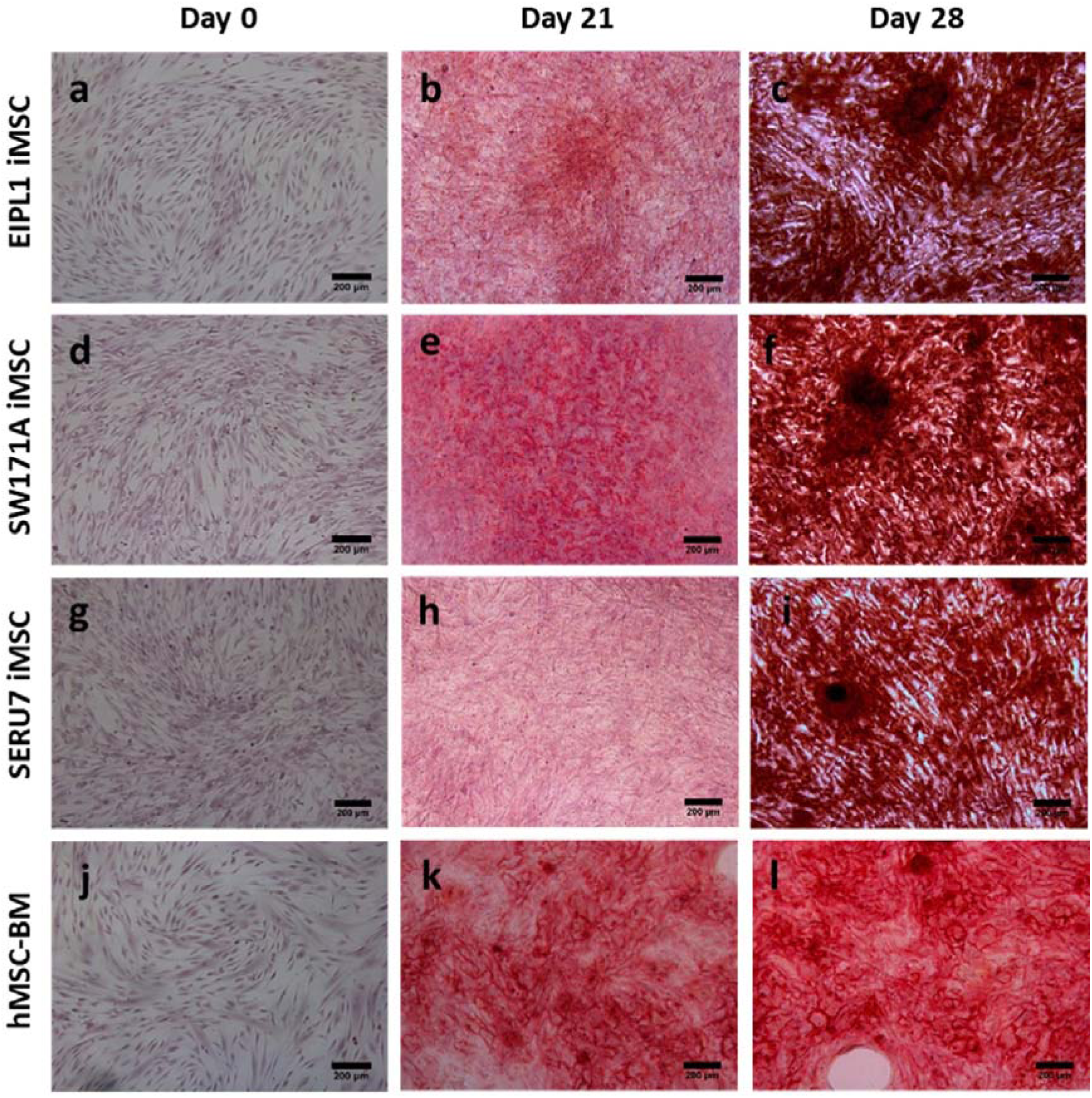
Characterisation of iMSC and hBM-MSCs via alizarin red staining. Both iMSCs and hBM-MSCs underwent osteogenic differentiation with noticeable mineralisation nodules being produced by the end of differentiation (day 28) as displayed by alizarin red staining. Images were collected on an Olympus IX83 inverted microscope and captured through MMI CellTools software at 10x magnification. Scale bars shown at 200 μm.

### Differentiation of iMSC into VSMCs

To produce VSMCs from iMSCs (the iMSC-VSMCs), we optimised a protocol where TGF-β1 and PDGF-BB induced iMSCs to form VSMCs in nine days as illustrated in Figure 3A. The hBM-MSCs ran in parallel with the iMSCs as controls. During the differentiation, VSMC-specific maker genes including *ACTA2, CNN1, TAGLN* and *MYH11* were gradually increased and reached a highly significant level by day 6 (Figure 3B). On day 9, the expression of these marker genes increased by around 5 to 50-folds between the iMSC-VSMCs and hBM-MSC-VSMCs depending on specific gene, comparing to that of the MSCs on day 0 (Figure 3B). Immunofluorescent staining showed a homogenous population of cells positive for VSMC-specific makers α-SMA, calponin, and SM22α in both iMSC-VSMCs and hBM-MSC-VSMCs by day 9 as compared to the undifferentiated iMSCs and hBM-MSCs (day 0) (Figure 3C). Both iMSC-VSMCs and hBM-MSC-VSMCs were positive for the calcium channel, voltage-dependent, L type, alpha 1C subunit (Cav1.2) (Figure 3C) that is usually highly expressed in the contractile VSMCs. The expressions of most VSMC marker genes did not further increase after day 9 (Figure 3B), therefore, the cells were switched to the VSMC maintenance medium at this time point and the expressions of VSMC marker genes were maintained on day 12 (Figure 3B). As anticipated, the expression of the pluripotent markers *Oct 3/4* and *SOX2* in the iMSC-VSMCs was significantly downregulated when compared to the undifferentiated iPSCs (**Figure S4**).

**Figure 3.**
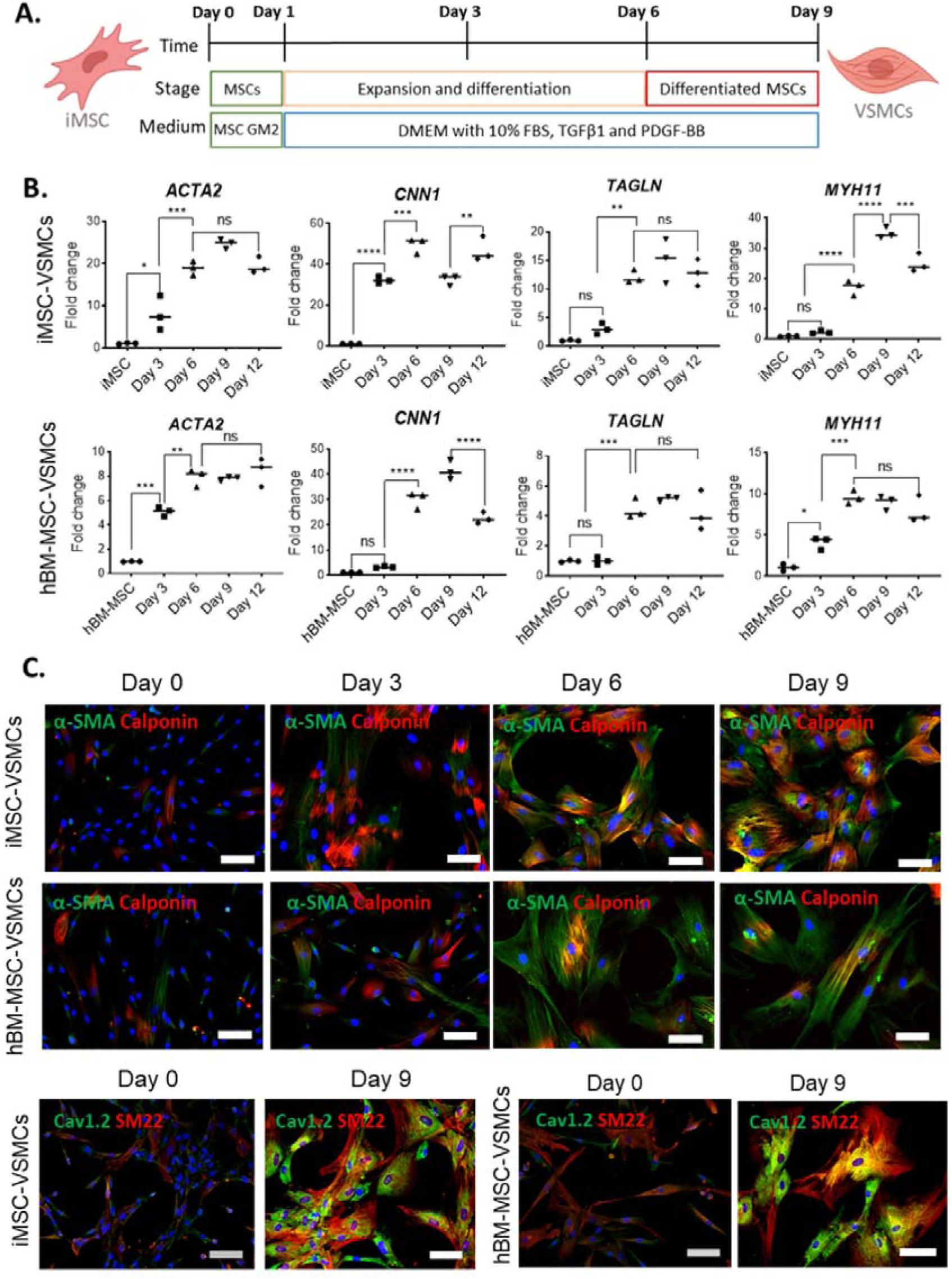
Detection of common VSMC markers in hBM-MSC-VSMCs and iMSC-VSMCs. A schematic of the VSMC differentiation protocol is shown (**A**). Fold gene expression increases in VSMC marker-genes α*-SMA*, *CNN1*, *SM22* and *MYH11* was also noticed throughout differentiation of hBM-MSC-VSMCs and iMSC-VSMCs (**B**) More homogenous population positive for contractile VSMC markers α-SMA and CACNA1C (green) as well as CNN1 and SM22 (red) could be observed by day 9 of differentiation via Immunofluorescence staining in hBM-MSC-VSMCs (C) and iMSC-VSMCs (**C**). Scale bars shown at 100 μm. Data significance is presented as *p < 0.05, ** P ≤ 0.01 *** P ≤ 0.001 **** p ≤ 0.0001 (n = 3)

### VSMC Differentiation in Silk Fibroin Coated Porous Electrospun Vascular Scaffolds

We have previously reported the development of a silk fibroin coated highly porous electrospun poly(L-lactic acid) (PLLA) fibrous vascular scaffold that was compatible to grow rat aortic smooth muscle cells ^19^. The production method and morphology changes throughout modifications cand be observed in (Figure 4A**, 4B**). Changes fibre topography and mechanical properties was also further analysed in this study (**Figure S5**). The scaffolds in this study were served for in-scaffold differentiation of iMSC-VSMCs to create cell laden TEVGs. To assess its suitability to manufacture TEVGs, we first assessed the biocompatibility of the scaffold to the iMSCs and hBM-MSCs, and particularly evaluated the effect of the silk fibroin coating on cell growth. After 10 days cell culture, the silk fibroin coated porous scaffold significantly favoured cell proliferation as shown by cell counting and the cellular metabolic activity as measured using alamar blue (Figure 4C) for both iMSCs and hBM-MSCs. Fluorescence staining with phalloidin showed that the iMSCs and hBM-MSCs growing in the silk fibroin coated porous scaffolds appeared to have better morphologies with a more spreading feature than the prestine and porous-only Scaffolds (Figure 4D). Thus, the silk fibroin coated porous electrospun scaffolds were chosen for fabricating the cell-laden TEVGs.

**Figure 4.**
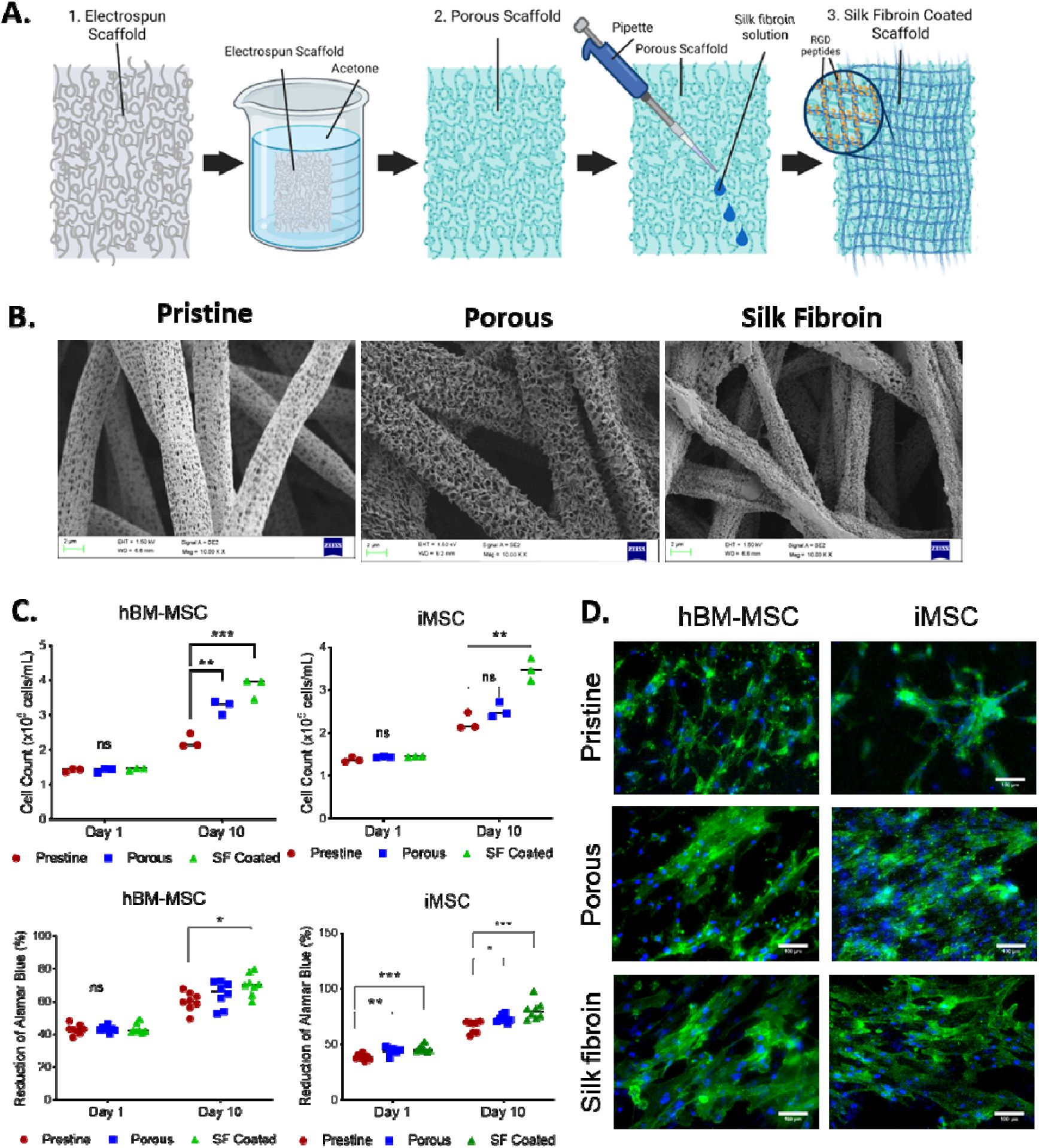
Proliferation and metabolic activity of hBM-MSC and iMSCs. A diagram of the modifications made to the pristine (1) by immersing in acetone ≥ 99.8% acetone to porous (2) fibres and covered with a 1% w/v silk fibroin solution to coat scaffolds (3)(**A**). Three different types of scaffolds were visualised under SEM imaging. Pristine scaffolds produced by electrospinning without any post-electrospinning modification (**B**). Cells count differences in iMSCs and hBM-MSCs seeded on pristine, porous and silk fibroin coated scaffolds after and reduction of AlamarBlue™ differences in iMSCs and hBM-MSCs seeded on pristine, porous and silk fibroin coated scaffolds after 10 days of culture (**C**) Density and morphological differences between the scaffold conditions were also observed via immunofluorescence staining with Phalloidin-iFluor 488 (A). Scale bars shown at 100 μm (**D**). Data significance is presented as *p < 0.05, ** P ≤ 0.01, *** P ≤ 0.001 (n =3).

Prior to the production of cell-laden TEVGs, In-scaffold differentiation of iMSCs and hBM-MSCs was carried out using the optimised method as mentioned above on 2.5 x 5.5 cm silk fibroin coated electrospun sheets. After 9 days, cell attachment was first assessed via phalloidin staining (Figure 5A) and differentiation of iMSC-VSMCs and hBM-MSC-VSMCs was assessed thereafter by positive staining of α-SMA and calponin (Figure 5B). Differentiated iMSC-VSMCs and hBM-MSC-VSMCs exhibited similar characteristic VSMC markers on the electrospun sheets coated with silk fibroin to those differentiated in the conventional tissue culture conditions without scaffold (Figure 3C).

**Figure 5.**
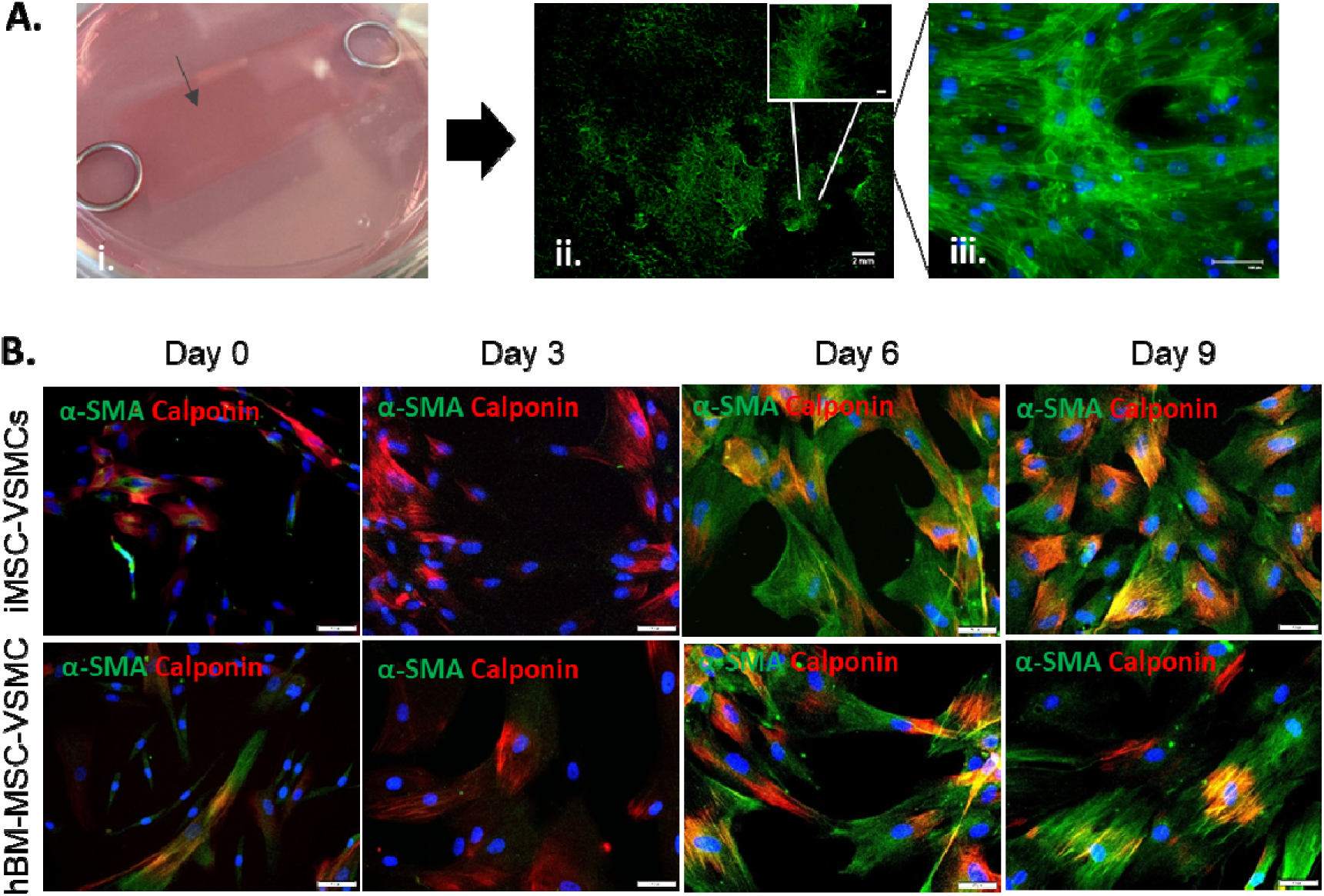
Detection of common VSMC markers in hBM-MSC-VSMC and iMSC-VSMC ladens electrospun scaffolds. Electrospun sheets coated with silk fibroin were seeded with hMSCs-BM (**Ai**). After 9 days, hMSCs-BM stained with phalloidin 488 showed to have adhered and spread throughout the scaffolds (**Aii and Aiii**). Scale bars shown at 2 mm for the wide view and 100 μm in the zoomed images. Images were aquired on an Olympus BX63 upright microscope using a using an DP80 camera (Olympus) through CellSens software. Homogenous population positive for contractile VSMC markers α-SMA (green) and CNN1 (red) could be observed in hBM-MSC-VSMCs and iMSC-VSMCs laden scaffolds after 9 days of differentiation. Scale bars shown at 50 μm at 20x magnification (**B**).

### Production and Characterisation of Tissue Engineered Blood Vessels

The overall approach to produce tissue engineered blood vessels in our study is illustrated in Figure 6A, where MSCs differentiated into VSMCs in a scaffold sheet that was then rolled into a tubular structure with a desired diameter, which was sealed with alginate (Figure 6A). Based on the in-scaffold VSMC differentiation parameters optimised, the iMSCs and hBM-MSCs were seeded on rectangular silk fibroin coated electrospun scaffold sheets (2.5 x 5.5 cm) mentioned above and differentiated for 9 days, as previously detailed, and then switch to the VSMC maintenance medium. Once maintenance media was added, VSMCs were further expanded over a 3-week culture period. The VSMC-laden sheet scaffolds were rolled on a 3 mm diameter stainless steel rod to generate multiple tubular shaped vessel constructs. The vessels were then dipped in a 3.5% alginate solution and cross-linked with calcium chloride to seal the constructs and maintain a vessel-like shape. The vessel mimics measured ∼3mm inner diameters, 0.51 mm wall thickness and ∼20 mm in length, similar to those of a coronary artery of similar calibre (Figure 6B). Longitudinal cross-sections were made, and immunofluorescence stained for VSMC markers. Images showed that both of the iMSC-VSMCs and hBM-MSC-VSMCs maintained their VSMC phenotype in the maintenance medium conditions as evident by the α-SMA and calponin positive cells observed across the length of the tube constructs (Figure 6C).

**Figure 6.**
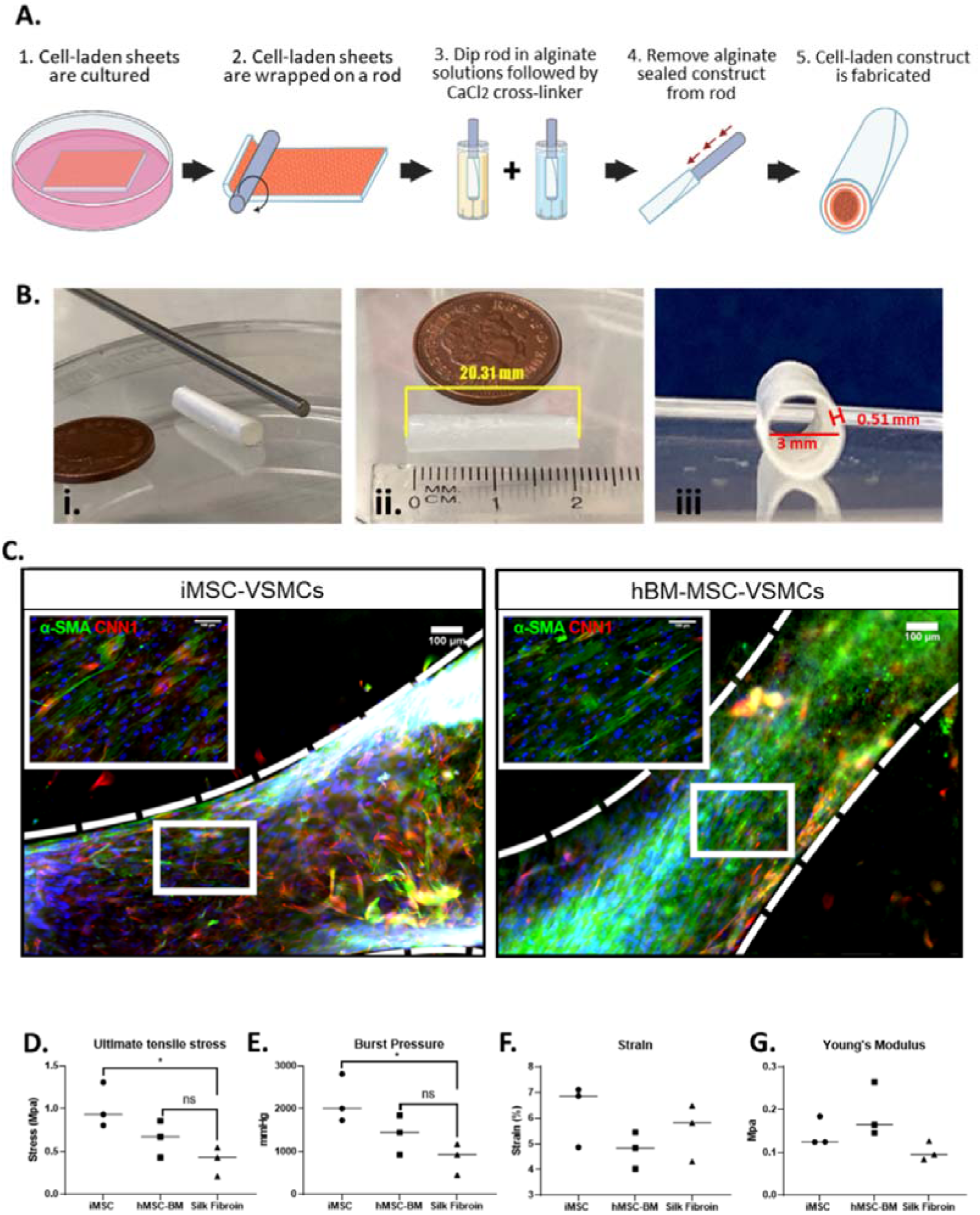
Vessel-like constructs derived from hBM-MSCs and iMSCs. Diagram of the fabrication process of tissue engineered blood vessels (**A**). The tube construct post-production can be seen next to the steel rod used to fabricate next and next to a penny for scale comparison (**Bi**). Length dimensions (**Bii**), wall thickness, and inner diameter dimensions (**Biii**) are also displayed. Immunofluorescence images of longitudinal cross-sections of tube constructs (B). Vessel mimics fabricated using both hBM-MSC-VSMCs and iMSC-VSMCs can be observed to be densely-populated with cells positive for α-SMA (green) and CNN1 (red). DAPI was counterstained to show nuclei. Scale bars shown at 100 μm. The representation of the scale is displayed in (**C**) Mechanical property differences in UTS (**D**), burst strength (**E**), young’s modulus (**F**) and strain (**G**) were also measured. Data significance is presented as *p < 0.05 (n =3).

The tissue engineered cell-laden vascular constructs were further assessed for their mechanical characteristics. Measurement of stress and strain mechanical properties showed that the hBM-MSC-VSMCs had a higher ultimate tensile stress (UTS) (0.655 ± 0.123 Mpa) and burst pressure (1408 ± 265.400 mmHg) compared to those from without cells (UTS 0.397 ± 0.098 Mpa, burst pressure 852.500 ± 211.800 mmHg) although no statistical differences were observed (Figure 6D**, 6E**). However more pronounced differences were observed in vessel mimics derived from iMSC-VSMCs with the UTS (1.017 ± 0.151 Mpa) and burst pressure (2184 ± 325.300 mmHg) significantly higher than the cell-free grafts (Figure 6D**, 6E**). Young’s modulus and strain were also measured which were similar between scaffolds with and without cells, although there is a trend of increase of the Young’s modules for the cell-laden grafts (Figure 6F**, 6G**). Overall, the mechanical properties of UST, burst pressure, young’s modules and strains of the tissue engineered grafts were all similar to autologous vascular grafts used in CABG to treat CAD ^10,27,28^.

## Discussion

In this study, we presented a new, rapid and simple method to fabricate tissue engineered blood vessels. The method demonstrated its capability to fabricate small diameter tubular constructs laden with in-scaffold production of VSMCs from iPSC-derived MSCs, and possessing native like mechanical properties. Thus, this strategy provided a potential approach to generate “off-the-shelf” TEVGs that are currently in urgent need due to scarcity of autologous donor vessels and poor synthetic alternatives currently available for CABG ^10^.

Various electrospun strategies have been applied to fabricate TEVGs with promising results *in vivo* including their performance in large animal models like sheep carotid arterial models ^17,29,30^. However, currently no electrospun TEVG has reached the clinic for CABG. The majority of the TEVGs reported to date were either not cell-laden but relied on host-cell infiltration post-implantation to remodel the grafts, or pre-seeded primary vascular cells during the fabrication process prior to implantation ^16,31^. This have been the approaches used for all TEVGs that have reached the clinic at present to treat occluded small-diameter vessels ^12,13,32,33^. However, these methods are confronted with significant downsides. Most noticeably, the use of primary cells requires an autologous source to avoid immunogenicity. The donor age or disease can significantly affect the functionality including the proliferation or remodelling ability of harvested cells. Primary cells usually have significantly limited lifespan *in vitro* as they become rapidly senescent ^21,34,35^. Stem cells have greater regenerative potential due to their infinite renewal capacity and the ability to differentiate into various cell types upon environmental stimuli, which provides an excellent source for fabricating TEVGs ^21,36,37^. MSCs are particular appealing among other stem cell types, like ESCs or iPSCs, despite limited plasticity, as they have immunosuppressive features which been extensively investigated clinically and have shown to prevent immunogenic responses upon implantation ^23,38–40^. Although iPSCs have remarkable potential to generate various types of tissue due to their plasticity, they lack the same immunosuppressive features and undifferentiated iPSCs have significant tumorigenic potential upon implantation ^41^. Thus, scaffolds used to bioengineer blood vessels in this study were seeded with hBM-MSCs or hiPSCs that were first derived in iMSCs prior to seeding or differentiating into VSMCs to minimize the risk of tumorigenicity.

VSMCs, as the most abundant cell type in arteries, play a critical role in providing mechanical support and controlling the constant circulation of blood flow in vasculature ^42,43^. Their contractile functions are key to the regulation of the vascular tone of blood vessels by buffering the pulsatile flow and maintain blood circulation ^44,45^. Due to their important roles, various tissue engineering strategies have focused in mimicking the middle layer (Tunica media) of native vasculature when fabricating blood vessels ^46–48^. In this study, we demonstrated a viable method to derive VSMCs from hiPSCs via a MSC intermediate. A monolayer defemination method was used to produce VSMCs in part to provide a more reproducible protocol with well-defined chemical factors to direct it towards an MSC intermediate stage. SB431542, a TGF-β1 pathway inhibitor, was used to induces an EMT-like process and an MSC maintenance media was later introduced to promote maturation of these cells towards and MSC phenotype. This strategy offers significant clinical advantages as it couples key beneficial features of both MSCs and iPSCs. One of these features are their ability of MSCs to prevent immunogenic responses by host tissue post-implantation which have also been displayed by iMSCs. In various studies, the expression of MHC class I, TLRs and PD-L1 in MSCs has demonstrated to aid in escaping attacks from activated natural killers and avoid an immunogenic response ^38,40,49–51^. This particular characteristic can enable grafts generated from MSC intermediates to have an allogenic clinical approach apart from the general patient-specific approach iPSCs can provide. Furthermore, MSCs have been also shown to secrete TGF-β1 and hepatocyte growth factor (HGF) in response to inflammation to inhibit the activation and functions of T Cells which can further reduce the chances of immunogenic responses ^23,51–53^. Therefore, although a patient-specific approach is viable using patients own reprogramed iPSCs, our approach also enables a potential for an “off-the-shelf” TEVG application using these key immunosuppressive characteristics MSCs possess by going through that intermediate stage. Moreover, Previous studies have shown that hiPSC derived MSCs could not only generate cells with a rejuvenated phenotype compared to adult MSCs but also enhance vascular regeneration and healing ^54–56^. Our derived iMSCs had yields of 50-89% depending on the MSC-specific marker (CD73, CD105 and CD90) without requiring a cell-sorting step and were largely absent for phenotypical MSC negative markers (≤18% positive cells). These measurements were comparable to those in hBM-MSCs and iMSCs were also able to undergo osteogenic differentiation like hBM-MSCs. While yield of subsequently derived VSMCs was not measured, VSMC marker genes significantly increased by the end of differentiation with homogenous populations positive for VSMC markers observed via immunostaining.

In a previous study, we assessed the biocompatibility of our electrospun scaffolds on A7r5 rat smooth muscle embryonic aorta cells ^19^. In this study we aimed to further validate these results with human cells and assess their biocompatibility over an extended culture period. Moreover, this study evaluated the suitability of these scaffolds to generate vascular tissue like constructs derived from stem cells. Our previous study demonstrated that cells laden on all electrospun scaffold conditions could support proliferation from day 1 to 4. Our present results with hBM-MSCs and iMSCs showed similar outcomes and up to 10 days. However, a clear improvement in proliferation was observed in cell-laden porous scaffolds compared to pristine scaffolds and further enhanced by the silk fibroin coating due to it well known RGD binding sites present. Albeit cell dependent, porous or rough surfaces have shown to improve cell attachment, proliferation, migration and viability as observed in this study ^57,58^. Silk fibroin, which demonstrated to also enhance cell function in this study, is a widely available biomaterial derived from silk and unlike other natural based biomaterials with mammalian cell adhesive motifs like collagen, gelatin or fibrin, it possesses remarkable mechanical strength. As a result, it has been extensively used for clinical applications such as biodegradable suturing materials or surgical meshes for wound healing ^59–61^. Therefore, due to their superior biocompatibility, silk fibroin coated scaffolds were used for further hBM-MSCs/iMSC to VSMC differentiation experiments and fabrication of tissue engineered blood vessels.

Like those cultured in conventional tissue culture conditions, iMSCs and hBM-MSCs were able to successfully differentiate in VSMCs when seeded on the elctrospun silk fibroin coated scaffolds. As a result, they were used to fabricate blood vessel mimics using sheet wrapping method. One common method used to fabricate tube scaffolds via electrospinning is to spin fibres onto a conductive mandrel that acts as a subtractive mould. When the mandrel is removed, a tube-shaped fibrous mesh is left ^62,63^. However, other methods like those that use electrospinning to fabricate fibrous sheet-like meshes and wrap them around a rod also exist. While both are viable methods, sheet-based strategies may be advantageous due to its simplicity and ability to generate multiple concentric layers like the tunica media in vessels as it wraps around the rod ^19,64–66^. Further, it may also be advantageous as the calibre of tube constructs can be easily adjusted during the fabrication process to best match comply with anastomosed vasculature if used for bypass for an “off-the-shelf” product. This was demonstrated during the optimisation of the fabrication process of the bioegineered blood vessels in this study (**Figure S6**). The bioengineered blood vessels in this study not only showed to possess tunic-medial like features with densely populated iMSC-VSMCs or hBM-MSC-VSMCs but those derived from iMSC-VSMCs had similar mechanical properties to common autografts used in CABG. Moreover, the burst pressure calculated for these blood vessels (2184 ± 325.300 mmHg) was significantly greater than similar strategies using hiPSC-VSMC derived electrospun blood vessels like those by Sundaram et al. ^37^ (600-800 mmHg) which are well below those of autologous vessels used for CABG. While this study successfully demonstrates a viable method to fabricate TEVGs with mechanical properties comparable to autologous grafts, limitations should be acknowledged.

The study was conducted entirely *in vitro*, and no *in vivo* testing was performed. Although the TEVGs showed promising structural, mechanical and biocompatible characteristics, their long-term performance, integration, and remodelling capabilities in a physiological environment remain to be evaluated. Nevertheless, based on the robust mechanical properties and biological characterisation outcomes observed, we believe the TEVGs developed in this study are ready for *in vivo* testing in animal models to assess their functional performance in a more clinically relevant setting. Future studies should focus on validating these constructs in preclinical models to further determine their suitability for clinical translation.

In conclusion, a viable and versatile method to fabricate tissue engineered blood vessels derived from different stem cell types was developed in the present study. A protocol successful at generating hiPSCs derived MSCs and subsequent iMSC derived MSCs was reported in this study that could also be applied to derive VSMCs for hBM-MSCs. Derivation of VSMCs from iMSCs and hBM-MSCs was also shown in electrospun scaffolds coated with silk fibroin to demonstrate stem cell derived vascular constructs could be generated. These vascular constructs included a bioegineered blood vessel with tunica-media like features after 3 weeks of culture. The VSMC-laden tunica media-like bioengineered vessels also demonstrated to possess similar mechanical properties to those of common autografts used in CABG making it a potential candidate for TEVG application although further evaluation *in vivo* would be required to assess its suitability. Lastly, our simple yet versatile approach has shown it has the potential to fabricate “off-the-shelf TEVGs that can potentially be used universally as allogenic sources due to immunoregulatory appeal MSCs possess or via a patient-specific approach using hiPSCs. Therefore, showing significant potential for clinical application.

## Author Contributions

The experiments in this study were performed by Luis Larrea Murillo, Zhongda Chen, Jun Song, Adam Mitchel and Steven Woods. Luis Larrea Murillo wrote the original draft of this research article, which was reviewed and edited with the help of all authors, especially Tao Wang. Luis Larrea Murillo prepared the figures, which were edited by Tao Wang. Luis Larrea Murillo, Tao Wang, Yi Li, Jiashen Li contributed in various aspects of study conceptualisation and formal analysis. All authors have read and agreed to the published version of the manuscript.

## Acknowledgements

The Author would like to gratefully acknowledge Dr Peter March for his help in the bioimaging facilities at the University of Manchester and Michael Jackson at the University of Manchester core facilities for his contribution on training and using the flow cytometry equipment.

