## Supplemental figures S1-S6 for "Blood Vessels Bioengineered from Induced Pluripotent Stem Cell Derived Mesenchymal Stem Cells and Functional Scaffolds"

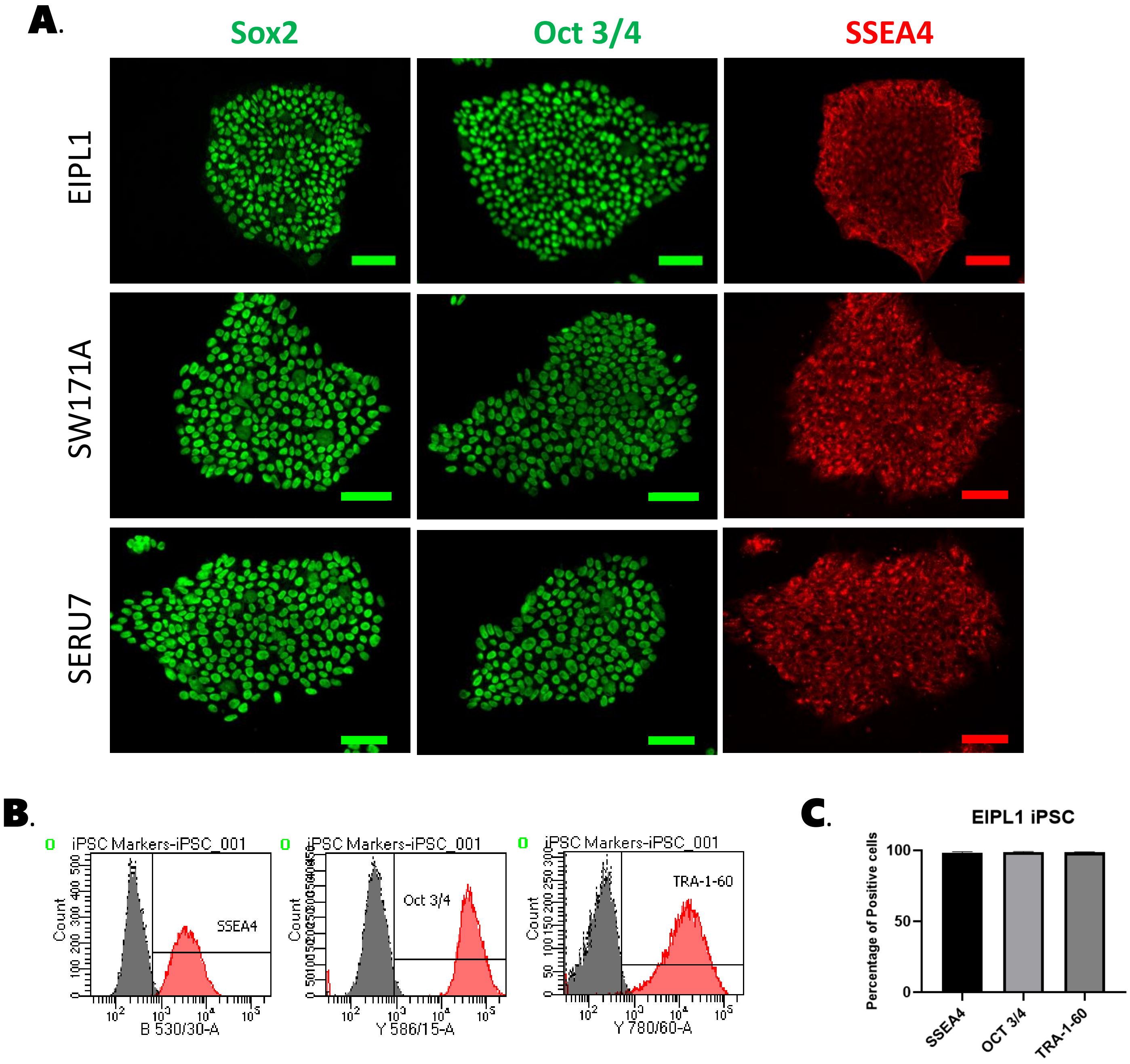


**Figure S1** Immunofluorescence staining of three iPSC lines (EIPL1, SW171A and SERU7) for the membrane pluripotency marker (SSEA4, red) and nuclear pluripotency markers (Oct 3/4 and Sox2, green). Scale bars shown at 100 µm. (**A**). Flow cytometry analysis of a representative iPSC line (EIPL1) for membrane (SSEA4 and TRA-1-60) and nuclear (Oct 3/4) pluripotency markers can also be observed by flow cytometry (**B**). EIPL1 hiPSCs were ≥98% positive for all three markers analysed via flow cytometry (n = 3) (C).

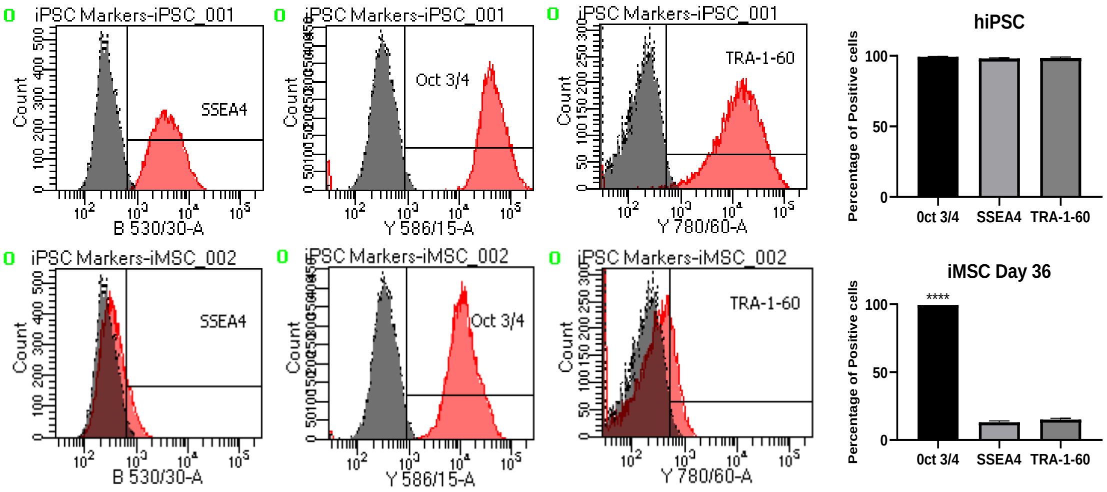


**Figure S2** When compared to hiPSCs via flowcytometry, the decrease of the percentage of cells positive for pluripotent markers SSEA4 and TRA-1-60 (≤15%) was observed with the exception of Oct 3/4 (98%) which remained unchanged once differentiation was concluded.

### Negative MSC Markers

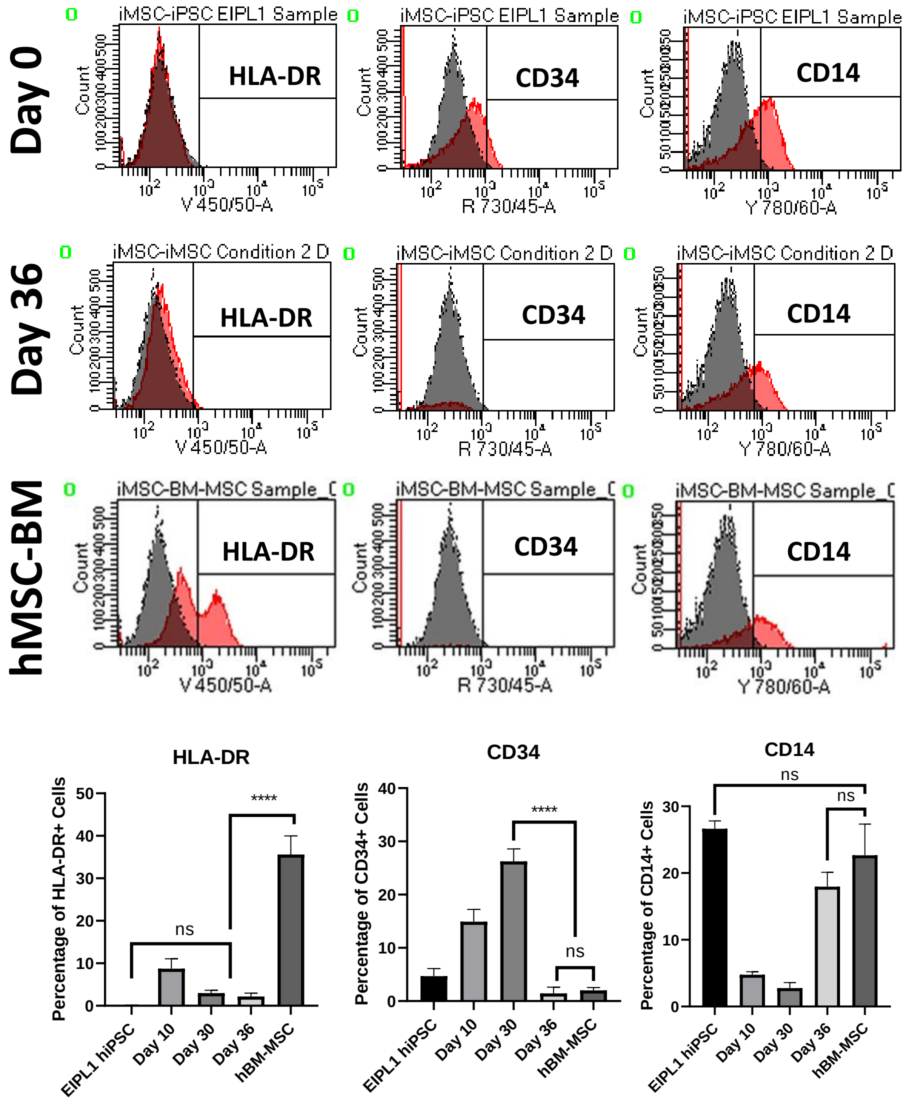


**Figure S3** Percentage of cells positive for HLA-DR, CD34 and CD14 decreased throughout iMSC differentiation although Percentage of cells positive for HLA-DR remained high in hBM-MSC.

| Target Gene | Primer Sequence |
|---|---|
| OCT 3/4 | Forward: CGAGAGGATTTTGAGGCTGC |
| | Reverse: CGAGGAGTACAGTGCAGTGA |
| SOX2 | Forward: GGAGCTTTGCAGGAAGTTTG |
| | Reverse: GCAAGAAGCCTCTCCTTGAA |

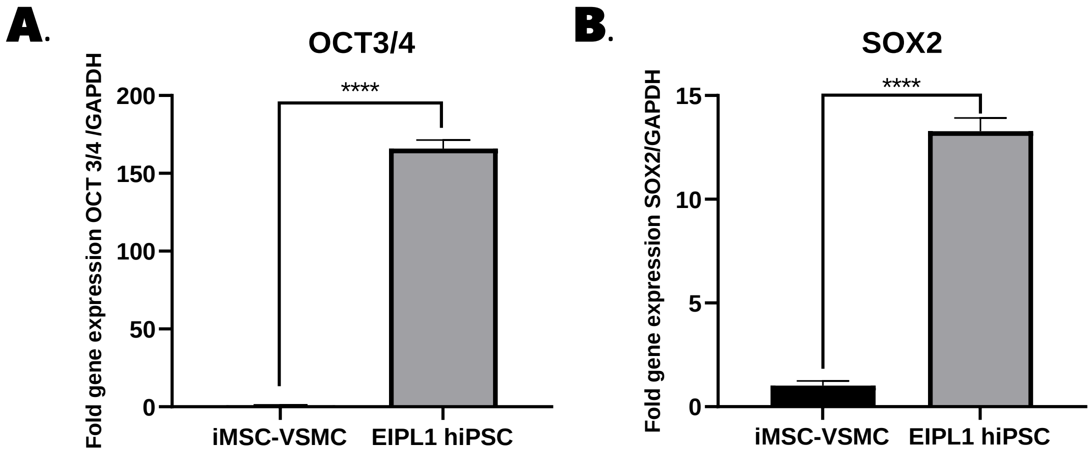


**Figure S4** Overexpression of pluripotent marker genes α-OCT 3/4 (A) and SOX2 (B) was observed in hiPSCs when comared to fully differentiated VSMCs from the same hiPSC cell line (EIPL1) Overexpression of these genes were from a 13.3 fold to as high as a 165 fold difference in hiPSCs. Data is presented as mean ± SEM, Unpaired t-tests, **** $P \leq 0.0001$, n=3

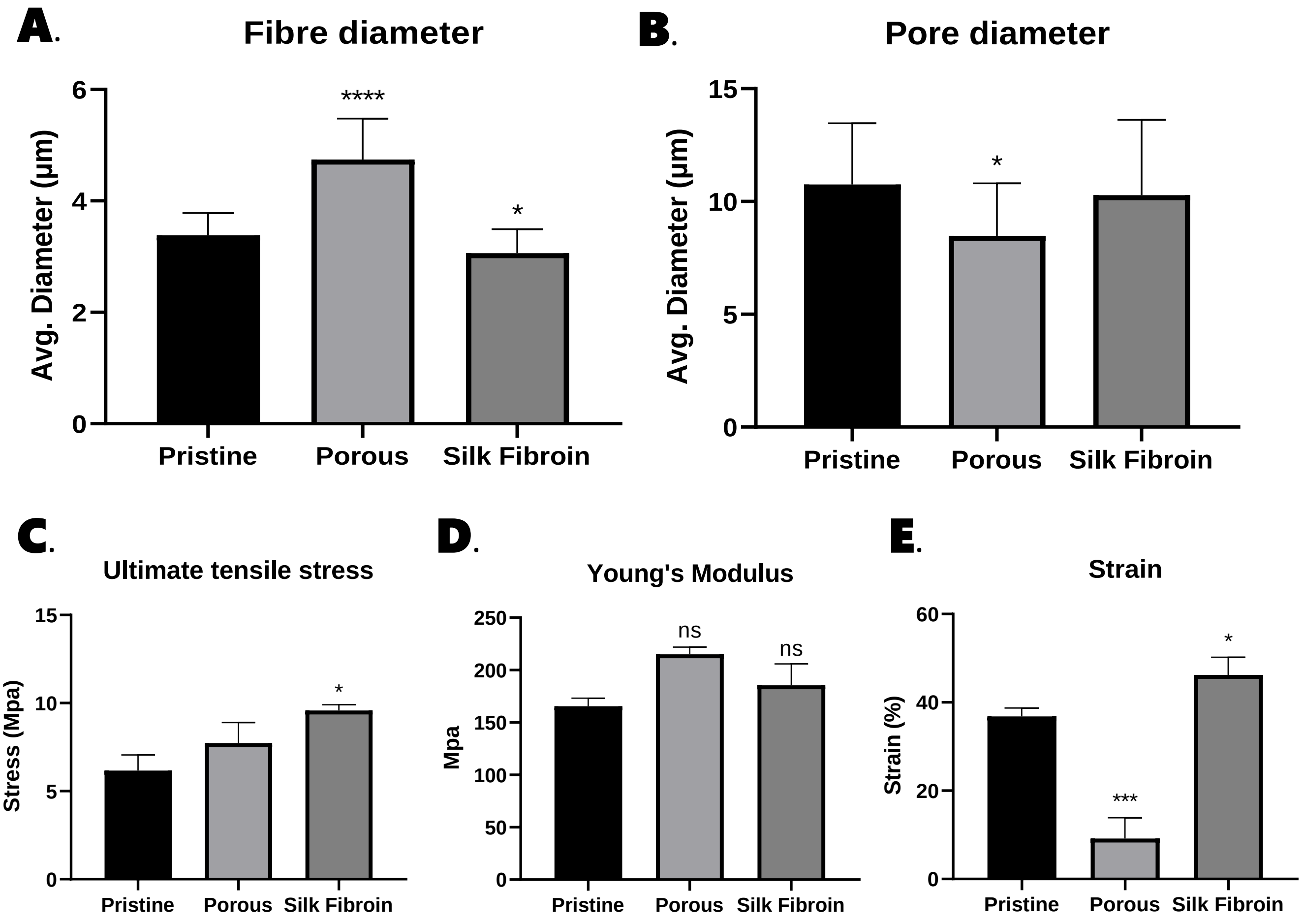


**Figure S5** Three different types of scaffolds were assessed for differences in typography, strength, deformation and stiffness. Differences Average fibre diameter (A). Average pore diameter (B) were observed. Ultimate tensile stress differences between the three scaffold conditions (C). Strain differences between the three scaffold conditions (D) Young's modulus differences between the three scaffold conditions (E).Average fibre diameter (E). Average pore diameter (F). Data are presented as n = 4 One-way ANOVA, *p < 0.05, **** p ≤ 0.0001.

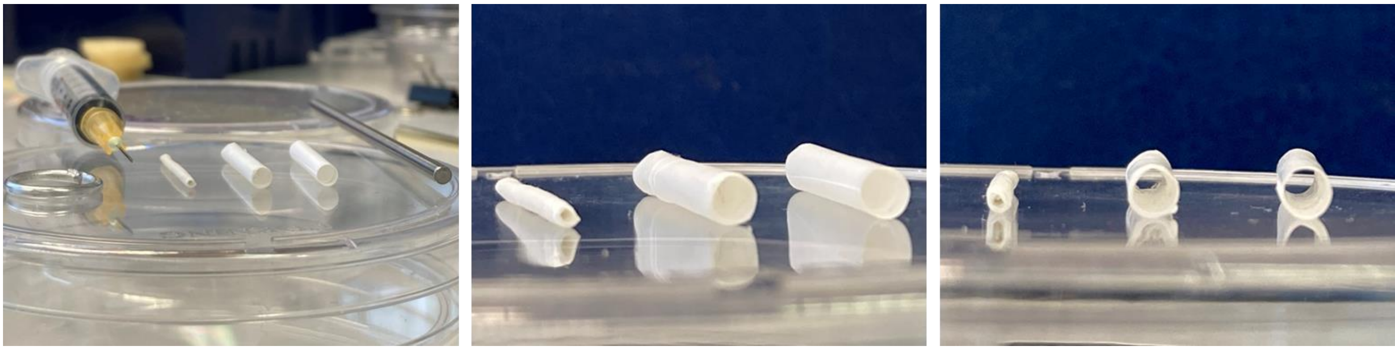

**Figure S6** Using the fabrication method above, cell-free tubular scaffolds were fabricated using 25-gauge syringe needles and 2.5 mm diameter stainless steel rods. Although these tubular constructs were not measured in detail to know their exact dimensions, the flexibility of this method demonstrates that the calibre of bioengineered vessels can be adjusted to best fit anastomosed vessels for bypass. The tube construct post-production can be seen next to a 2.5 mm steel rod and a 25-gauge syringe needle used to fabricate tube constructs next and next to a jump ring for scale comparison on the left. The images on the middle and right show zoomed in images of the constructs at different angles.
